# Immediate neural network impact after the loss of a semantic hub

**DOI:** 10.1101/2022.04.15.488388

**Authors:** Zsuzsanna Kocsis, Rick L. Jenison, Thomas E. Cope, Peter N. Taylor, Ryan M. Calmus, Bob McMurray, Ariane E. Rhone, McCall E. Sarrett, Yukiko Kikuchi, Phillip E. Gander, Joel I. Berger, Christopher K. Kovach, Inyong Choi, Jeremy D. Greenlee, Hiroto Kawasaki, Timothy D. Griffiths, Matthew A. Howard, Christopher I. Petkov

## Abstract

The human brain extracts meaning from the world using an extensive neural system for semantic knowledge. Whether such broadly distributed systems^1–3^ crucially depend on or can compensate for the loss of one of their highly interconnected hubs^4–6^ is controversial^4^. The strongest level of causal evidence for the role of a brain hub is to evaluate its acute network-level impact following disconnection and any rapid functional compensation that ensues. We report rare neurophysiological data from two patients who underwent awake intracranial recordings during a speech prediction task immediately before and after neurosurgical treatment that required disconnection of the left anterior temporal lobe (ATL), a crucial hub for semantic knowledge^4–6^. Informed by a predictive coding framework, we tested three sets of hypotheses including *diaschisis* causing disruption in interconnected sites^7^ and *incomplete* or *complete compensation* by other language-critical and speech processing sites^8–10^. Immediately after ATL disconnection, we observed highly specific neurophysiological alterations in the recorded fronto-temporal network, including abnormally magnified high gamma responses to the speech sounds in auditory cortex. We also observed evidence for rapid compensation, seen as focal increases in effective connectivity involving language-critical sites in the inferior frontal gyrus and speech processing sites in auditory cortex. However, compensation was incomplete, in part because after ATL disconnection speech prediction signals were depleted in auditory cortex. This study provides direct causal evidence for a semantic hub in the human brain and shows striking neural impact and a rapid attempt at compensation in a neural network after the loss of one of its hubs.

## MAIN TEXT

Speech is an important acoustic signal used by humans to communicate meaning. Successful comprehension of speech sounds engages a broad neural network in the human brain that rapidly decodes complex acoustic signals and binds them with categorical and contextual information crucial for concept formation and semantic knowledge^9,11–13^. By *distributed coding* accounts, this neural system is extensive^1,2^. For instance, decoded representations can be observed as semantic ‘tiles’ throughout the human brain^1,2^, including in speech processing temporal lobe regions and language-critical sites for semantic and syntactic functions in the inferior frontal gyrus (IFG)^8^. A *hub-and-spokes* model posits that these distributed semantic representations crucially rely on the anterior temporal lobe (ATL) as an indispensable hub^4,6^ that mediates learned associations distributed throughout cortex^14^. The ATL is highly interconnected and its hub-like functions both establish semantic representations elsewhere and send efferent predictions about forthcoming speech signals via interactions with the other nodes or ‘spokes’ in the network^15,16^. By this model, the loss of the ATL would severely disrupt semantic processes. The model is supported by patient studies, whereby stroke, surgical or degenerative damage to the ATL impairs responsive naming^17–19^, semantic memory^20–23^, and reduces the efficiency of language processing^24,25^. However, a direct causal test of the functional relationship between the hub and the distributed spokes has been outstanding^4^, and it remained unknown what the immediate site-specific neurophysiological impact and potential compensation on a human neural network might be after the loss of one of its hubs.

In this study, we directly compared pre- and post-disconnection intracranial recordings after a rare set of neurosurgical treatment procedures carried out in two patients. These data are rare because both patients participated in an acute, one-day left hemisphere ATL disconnection procedure to access and remove a temporal lobe seizure focus. Moreover, both patients had similar pre- and post-disconnection electrode coverage for clinical monitoring that included speech responsive IFG and auditory cortical sites (Fig. 1A-D), and they were tested awake during both recording periods as part of clinical language mapping. By comparison, most human neurosurgery patient studies provide intracranial site-specific neurophysiological information prior to, but not after the surgical treatment, or perioperative knowledge stems primarily from noninvasive neuroimaging. Although animal models provide precision in studying hub-like connectivity and function at cellular levels^26^, humans are the ideal model for speech and language semantic functions. Guided by three sets of hypotheses framed within a predictive coding framework^27–29^, this study provides direct causal evidence that implicates the ATL as a key hub for speech semantic processes and reveals remarkably rapid, albeit incomplete, neurophysiological attempts at compensation by a neural network after losing a hub.

**Figure 1.**
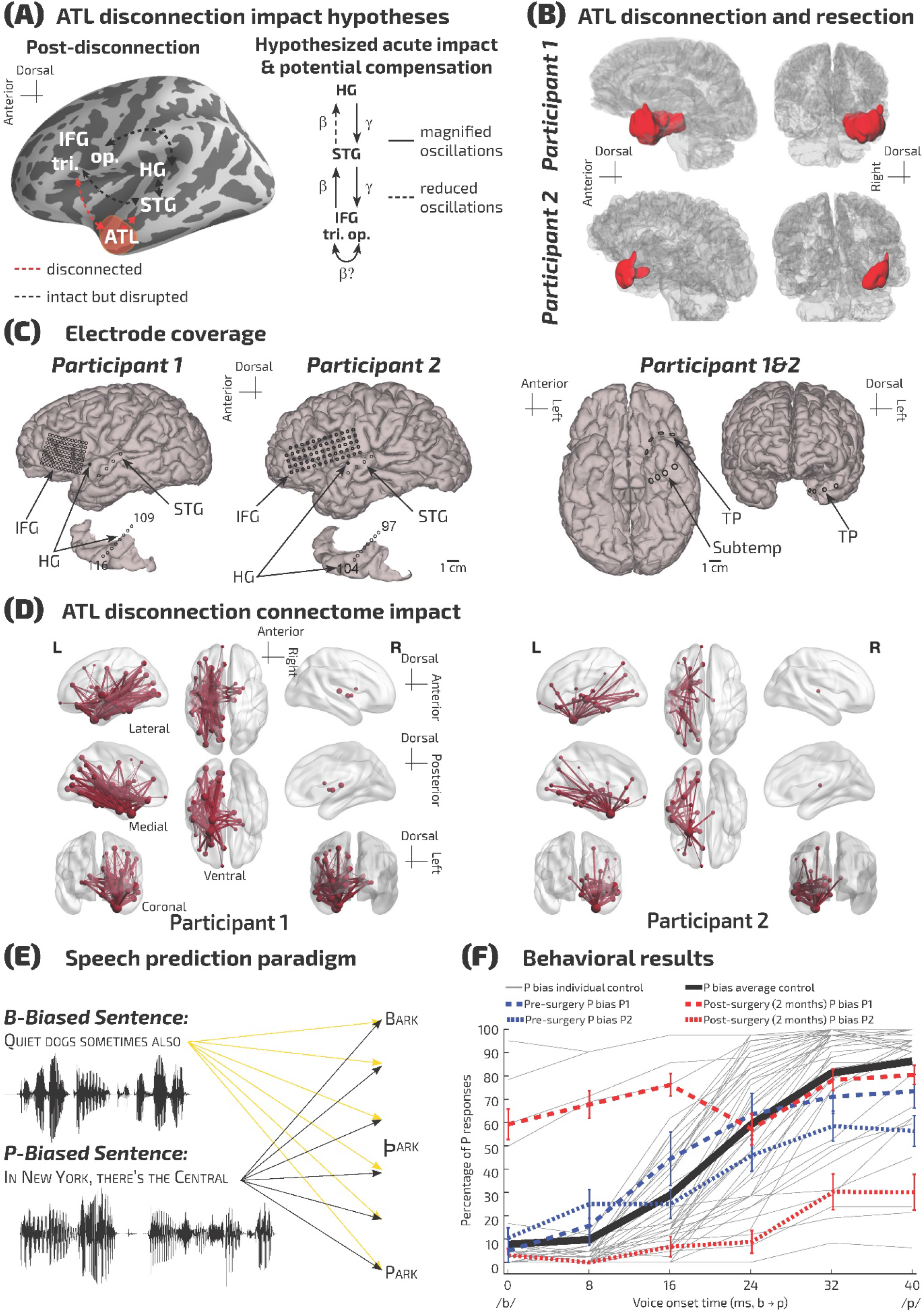
Participant electrode coverage, ATL disconnection structural impact, task and behavior. (A) Illustrated hypotheses: left hemisphere ATL disconnection to access the seizure focus is hypothesized to result in disruption and acute compensatory changes in neural signaling within IFG and auditory cortical subregions distant from the surgical resection site. (B) Reconstructed surfaces showing the disconnected regions for each participant in red; P1’s region also includes the MTL resection, which occurred after the ATL disconnection and recordings reported here (Suppl. Fig. 2). (C) Electrode coverage in the two participants. (D) Impact of ATL disconnection on structural connectivity pathways: Diffusion-weighted imaging derived networks show the reduction of streamlines caused by the ATL disconnection. Tube thickness is proportional to the percentage of streamlines removed from that connection. The size of each sphere represents the percentage reduction in the number of streamlines connecting that region. (E) Depiction of the speech prediction paradigm. Participants listened to /b/ or /p/ word biasing sentences with the final target word manipulated along a /b/ to /p/ voice onset time (VOT) continuum. (F) Behavioral responses under the /p/ bias condition, as a function of the /b/ to /p/ VOT continuum and the percentage of P responses given by the participants. The thin gray lines show each control participant (individuals without epilepsy who performed the task, based on ^37^) and the thick black line their average psychometric function. The blue dashed lines show the responses of P1 and P2, which fit within the normative distribution established by the control participants. The red dashed lines show the patients’ results 2 months after their ATL disconnection surgery, showing substantial disruption of their psychometric functions.

### Characterizing acute impact and levels of compensation

The possible acute impact on the fronto-temporal system following the loss of the ATL can be characterized along three sets of hypotheses. 1) *Diaschisis* stipulates disruption and neurophysiological impact on distant ATL interconnected sites^7^, including speech responsive inferior frontal and auditory cortical regions. A diaschisis hypothesis goes beyond the expected immediate impact in areas near the surgical site, predicting disruption at distant but interconnected fronto-temporal sites. Full support for a diaschisis hypothesis would demonstrate an irreplaceable role of the ATL in this acute phase. 2) *Incomplete immediate compensation* could result in, amidst disrupted functions, *increased* functional interconnectivity^10,30^ between frontal and auditory cortical sites involved in semantic and speech processes^15,16^. Support for this hypothesis would both underscore the important role of the ATL and demonstrate the capability of a neural network for rapid neurophysiological compensation. 3) *Full immediate compensation* is possible via the remaining brain areas, including the contralateral ATL, together with the language critical IFG (pars triangularis and opercularis) and speech processing areas in auditory cortex. The intact areas could at least temporarily stabilize speech representations and predictions in the network. Support for this hypothesis is expected in the form of minimal to no change in speech semantic processes in the fronto-temporal network, provided that the ATL is not indispensable for these functions.

To test the hypotheses, we relied on the canonical microcircuit implementation of the predictive coding framework to generate more concrete predictions about both the form of disrupted and potentially compensatory site-specific neurophysiological responses^27–29^. First, the framework posits that higher-order areas, like the ATL which generate the relevant top-down predictions should relay feedback predictive signals to sensory cortex in the form of lower frequency oscillations (e.g., beta/alpha; < 30 Hz)^31–34^. Second, when speech predictions are affected, as might be expected by the loss of the ATL, sensory cortex should generate strong feedforward prediction error signals at higher frequencies (e.g., gamma; > 30 Hz)^27,32,35^. Third, higher-order language-critical areas such as IFG and auditory cortical sites capable of compensating for the loss of predictive signals from the ATL could increase their interconnectivity in an attempt to reinstate affected semantic prediction signals. The hypotheses are illustrated in Fig. 1A. We ensured that all main interpretations are based on significant and consistent effects in both participants. We obtained evidence for both the *diaschisis* and *incomplete immediate compensation* hypotheses, largely in the forms specified by the predictive coding framework.

### Post-acute phase behavioral impact on semantic perception

To evaluate the post-acute behavioral impact on speech predictions in the two patients, we employed an established task where higher-level sentence context influences speech perception and predictions^36,37^. The task was conducted with the two participants 2-6 weeks before and 2 months after the day of the surgical procedure. It consisted of listening to sentences that biased the perception of a final target word. The task allowed us to assess both congruent semantic predictions (e.g., “Quiet dogs sometimes also <u>bark</u>”; Fig. 1E) and incongruent conditions (e.g., “Quiet dogs sometimes also <u>park</u>”). The voice onset time (VOT) of the target word was manipulated along a perceptual continuum from /b/ as in ‘bark’ (short VOTs) to /p/ as in ‘park’ (long VOTs), with several different /b/ and /p/ word pairs being used. Thus, the preceding sentence context, alongside a simple VOT change in the first phoneme of otherwise acoustically matched target words biases the semantic perception of the words.

In the presurgical testing period, the two participants’ phoneme categorization functions displayed a large initial semantic context effect: both participants tended to respond having heard more /b/ words in the /p/ bias condition than controls (Fig. 1F), and they almost never responded having heard /p/ in the /b/ bias condition regardless of the VOT (Supplementary Fig. 1). Beyond this effect, their /p/ bias condition functions were well within the normative range (Fig. 1F), even as their /b/ bias condition performance was at floor (Suppl. Fig. 1). This initial effect is larger than the control participants potentially because the patients’ left ATL pathology for which they were being treated affected their ability to integrate semantic expectations and perceptual cues to some extent. With their pre-surgical performance as a baseline, two months after the surgery their /p/ bias psychometric functions showed even greater disruption, substantially deviating from their pre-operative state, while the /b/ bias condition results remained at floor. This was statistically confirmed by a logistic mixed model run separately for each patient, which showed a significant or near significant main effect of Disconnection for both patients (P1: *Z* = 3.44, *p* < 0.001; P2: *Z* = -1.95, *p* = 0.051), and a significant interaction of VOT and Disconnection for P1 (*p* = 0.0056). See Suppl. Table 1 for the complete results and Suppl. Fig. 1 for further details.

Both participants performed very well on the control tasks in which there was no lexical competition, and the semantic context was never incongruent with the VOT. Specifically, before and after the surgery, their performance was nearly perfect on filler trials where the target word was always consistent with the priming sentence (e.g., “This wall needs another coat of <u>paint</u>” has no /b/ word counterpart; pre-op: 100% correct performance in both participants; post-op: 98% for P1; 100% for P2). Their performance on catch trials where the target word was presented visually as a written word along with another word was also very high (pre-op: 94% for P1; 98% for P2; post-op: 92% for P1; 98% for P2). The overall behavioral results indicate that the surgical procedure exacerbated an impairment in the inability of both patients to integrate VOT information with semantic expectations, amidst otherwise intact attentional or general meaning-related abilities, as assessed by the control tasks.

During the recordings before and after the ATL disconnection procedure in the operating room, the participants were awake and listened to a shortened passive version of the speech priming task during the intracranial recordings. We ensured both patients were awake and engaged with the clinical and research teams during the two recording periods. Moreover, during the recordings, P2 took part in a simple tone detection task that was interleaved with the speech priming task to ensure that they were awake and engaged. Their tone detection performance was high before (100% correct, 24/24 trials) and after (88% correct, 21/24 trials) the ATL disconnection procedure.

### Neural system state prior to ATL disconnection

Diffusion MRI data showed that the ATL in the two patients was structurally highly interconnected with frontal and temporal areas (Fig. 1D; Suppl. Fig. 2). Pre-surgery, the ATL also exhibited functional hub-like properties (Fig. 2D). Figure 1C shows the electrode coverage, which includes left hemisphere IFG, superior temporal gyrus (STG) and Heschl’s Gyrus (HG). Temporal pole (TP) contacts within the ATL were only viable pre-disconnection. Node hubness can be quantified using graph theoretic metrics such as weighted *degree centrality*, defined as the total strength of links incident upon a specific node^38^. As a measure of node hubness, we calculated the sum of each node’s bidirectional (in and out) functional connectivity edge weights taken from the results of the effective connectivity analysis using state-space Conditional Granger Causality (CGC). The results show that before ATL disconnection the temporal pole contacts in this region are hub-like within the recorded network in beta and gamma frequencies (Fig. 2D; Suppl. Fig. 3).

**Figure 2.**
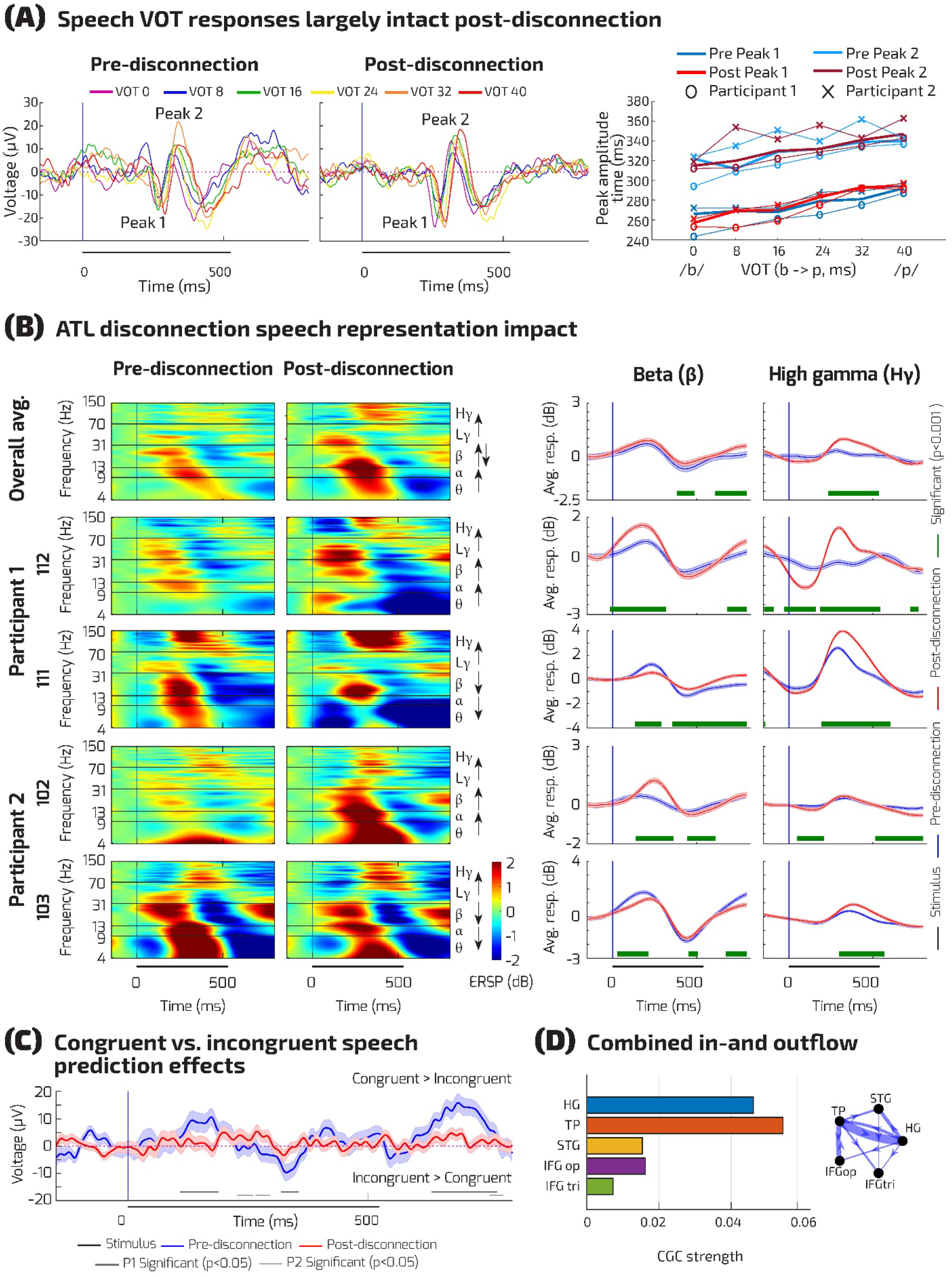
ATL disconnection impact on auditory cortical speech representations. (A) Posterior-medial Heschl’s Gyrus (HG) responses to the different VOTs in the speech sounds, averaged across the two participants and recording contacts in this region (N = 8). Onset of the target word is 0 on the horizontal axis. Right: VOT peak amplitudes for the participants, combined and shown separately. (B) Neurophysiological responses to the target words are selectively disrupted in HG. Left plots show the event-related spectral perturbations (ERSPs in dB) across the frequency bands (theta, alpha, beta, low and high gamma). Right shows a summary for beta (13-30 Hz) and high gamma (70-150 Hz; Suppl. Fig. 3 shows the other frequency bands). Top plots show speech representations averaged across contacts and the two participants. Bottom plots show the consistency of effects by participant and exemplar contacts. Right: blue lines show the pre-disconnection and red lines the post-disconnection responses (error bars are +/- Standard Error of the Mean (SEM) across trials). Statistically significant differences between the effects pre- and post-disconnection are shown in gray bars for each participant separately (cluster-based permutation test, *p* < 0.001). (C) Difference plots of the auditory evoked response in HG to the same speech sounds, contrasted by whether they were preceded by a congruent or incongruent predictive sentence. Blue shows the pre-disconnection congruency vs. incongruency mismatch response. This mismatch response is disrupted post-disconnection (red; cluster-based permutation test, *p* < 0.05 for at least 25 ms. (D) Gamma band Conditional Granger Causality measure of hubness showing the TP electrodes are functionally hub-like in beta and gamma.

**Figure 3.**
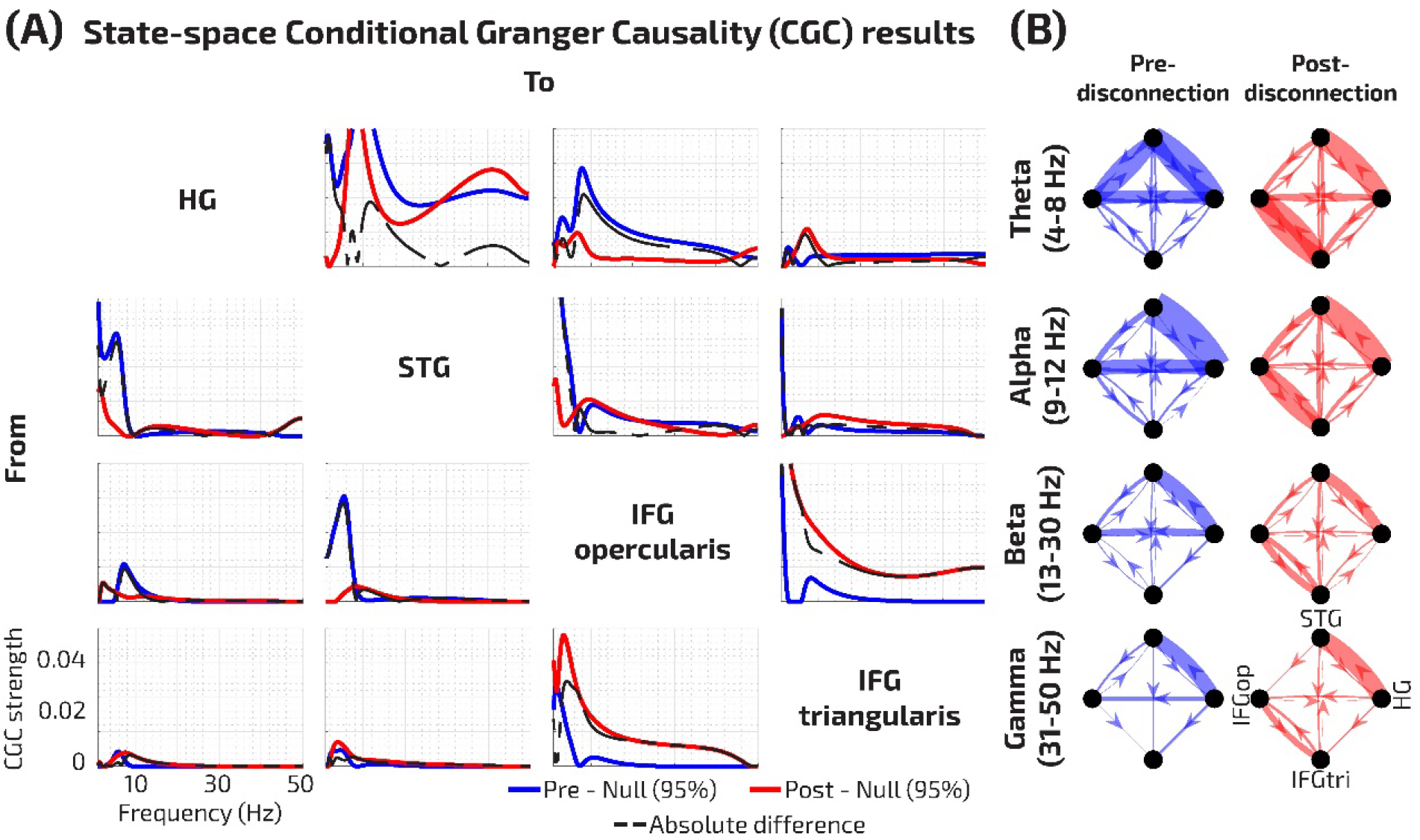
ATL disconnection impact on fronto-temporal network effective connectivity. (A) Combined frequency-resolved CGC spectral estimates calculated over 0 to 2 seconds relative to the onset of the target word, showing effective connectivity results between the recorded nodes in the fronto-temporal network (HG, STG, IFG p. opercularis and triangularis). Only statistically significant effects are shown for all spectra; the phase-randomized null distribution 95^th^ % was subtracted from each of the pre- (blue) and post-disconnection (red) conditions. A permutation test of spectral difference between pre- and post-disconnection was also performed (dashed black lines; one sided *p* < 0.05; Methods). Directions of influence from regions of interest (rows) to recipient regions (columns) are shown. (B) Strong dynamic directional influences are observed between all state-space modeled time-series in HG, STG and the IFG subregions, for each of the oscillatory frequency bands (theta, alpha, beta, gamma). The thickness of the directional edges represents the strength of the connection.

In both participants, we observed the expected strong broadband (3-150 Hz) oscillatory responses to the speech sounds in the auditory cortex (HG: Fig. 2; STG: Suppl. Fig. 4), with lower-frequency (< 30 Hz) speech responses in IFG including response suppression, as previously reported^39^ (Suppl. Fig. 4). Moreover, the auditory cortex showed clear speech VOT representations, seen as systematically time-shifted auditory cortical potentials to the VOTs pre-disconnection (Fig. 2A). Speech prediction-related neurophysiological responses in HG were also observed, including mid-latency (270-350 ms) evoked responses after target word onset that were stronger for incongruent compared to congruent word conditions (Fig 2C).

**Figure 4.**
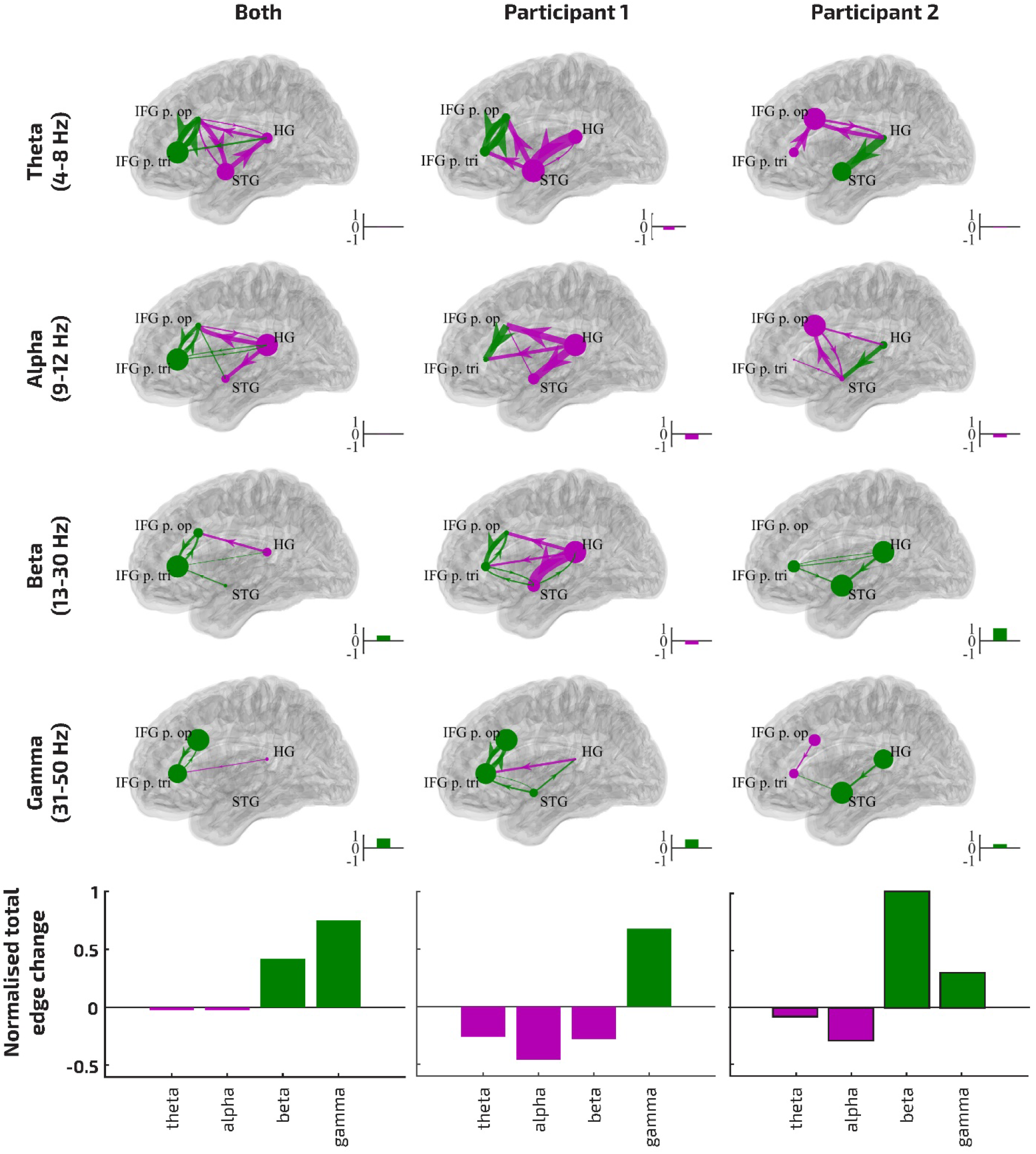
Effective connectivity alterations shown combined and by participant. Significant increases (green) or decreases (purple) post-disconnection in CGC edge weights (line thickness) and node hubness (inflow + outflow; node size/color) are depicted overlaid on a template brain. Results are shown for both participants combined and each participant separately (graphs, left to right), further separated by frequency bands (graphs, top to bottom). Additionally, for each plotted graph, total network increases/decreases in edge weight are shown normalized relative to total significant absolute edge weight changes (inset bars; also shown aggregated into summary plots, bottom). The normalized total network edge weight changes reveal that both participants exhibit relative increases in network-wide functional connectivity in the gamma band, and relative decreases in lower frequency bands. For congruency-related CGC effects see Suppl. Fig. 7.

### Neural network alterations after ATL disconnection

We first evaluated the structural connectome impact of the ATL disconnection procedure by comparing diffusion MRI data obtained 2-6 weeks before and 2 months after the surgical procedure. Structural connectivity analyses showed that the ATL disconnection affected interconnectivity with other frontal and temporal lobe areas via ventral pathways (Fig. 1D; Suppl. Fig. 2). Dorsal pathways between IFG and auditory cortex, including the arcuate fasciculus, were unaffected. Figure 1B shows the post-surgical lesion site, determined from pre- and post-surgical structural T1-weighted images which includes the ATL surgical procedure in both participants and the further MTL resection in P1. The cortical tissue affected *only* by the ATL disconnection procedure in both participants is shown in Suppl. Fig. 2. The surgical procedures and epilepsy pathologies were not identical in both participants, however their overall connectivity matrices before disconnection were highly similar (Spearman’s rho = 0.82; Suppl. Fig. 2B). The key difference in the surgical procedures was that P1 had more extensive temporal pole removal than P2. Additionally, P1’s ATL disconnection procedure also resulted in reduction of tissue in the superior and middle temporal cortical areas (Suppl. Fig. 2A). Supplementary Table 2 shows that the physiological parameters during the surgical procedure were comparable during the two recording periods. Anesthesia had been discontinued at least 20 minutes prior to the testing periods in both patients, with the ATL disconnection procedure completing ∼23 minutes prior to the commencement of the intracranial recordings reported here.

After ATL disconnection we obtained evidence for both unaffected and altered site-specific neurophysiological signals, including functional interactions between sites. One of the few neurophysiological signatures that remained largely unaffected was the auditory cortical responses to the VOTs pre- and post-disconnection in both participants (Fig. 2A; Wilcoxon signed-rank test respectively in Peak 1, P1: *Z* = 4.0, *p* = 0.375, P2: *Z* = 1.0, *p* = 0.125; Peak 2, P1: *Z* = 3.5, *p* = 0.375, P2: *Z* = 0, *p* = 0.0313). In support of the predictive coding framework hypotheses (Fig. 1A), in HG we observed strikingly magnified post-disconnection speech-related high gamma responses, significant in both participants (Fig. 2B). Relative to the pre-disconnection data, after the disconnection there was a magnified early suppression of high gamma power (∼0-250 ms) followed by later (∼250-500 ms) high gamma magnification to the target speech sounds (Fig. 2B right; cluster-based permutation test, *p* < 0.001). Lower frequency beta band responses were also affected post-disconnection, but differentially across HG recording contacts, with some responses enhanced and others attenuated across individual recording contacts (Fig. 2B). After ATL disconnection, theta and alpha band responses in HG yoked in directionality with effects across the recording contacts observed for beta. Higher-level stages in the auditory cortical hierarchy (STG) and the IFG pars triangularis showed mainly disruption of speech signal processing following ATL disconnection (Suppl. Fig. 4). Interestingly, the theta-based response in the IFG pars opercularis was significantly *increased* post-disconnection in both participants (Suppl. Fig. 4). Crucially, the neurophysiological semantic incongruency effect was depleted post-disconnection in HG (Fig. 2C; cluster-based permutation test, *p* < 0.05), and the later congruency-related components in IFG showed significant alteration post-disconnection (Suppl. Fig. 5) in both participants. This demonstrates that alterations in oscillatory fronto-temporal interactions were ineffective in rescuing neural speech-predictive responses to the target words. We also evaluated the neural response patterns to the speech sounds and the theta-gamma coupling response to speech, a known neurophysiological feature of fronto-temporal responses to speech^40^. There was an impact on the neural response pattern to speech that was consistent in both participants (Suppl. Fig. 6), however theta-gamma coupling was only evident in one of the participants pre-disconnection and is thus not considered further.

To identify whether and which post-disconnection effects resulted in altered directional functional connectivity, we relied on the spectrally-resolved state-space CGC analysis^40^. The CGC effects that significantly differed post-disconnection are shown combined and by participant in Figs. 3Ȧ4. Two observations are most striking. First, although there was a general disruption in effective connectivity, seen as significant *decreases* in CGC values post-disconnection (purple arrows, Fig. 4), certain interactions between the frontal and auditory cortical sites *increased* post-disconnection (green arrows, Fig. 4). Correspondingly in both participants, the hubness measure revealed increases in the centrality of selected nodes across the recorded fronto-temporal network, particularly in the gamma band (larger dots at each node, Fig. 4). Second, relative *decreases* in centrality dominated the network in the lower frequency bands in both participants (theta and alpha; purple dots, Fig. 4). Despite these commonalities, the spatial pattern of increased functional connectivity post-disconnection was mirrored across the frontal and temporal lobes in the two participants (pattern of green arrows, Fig. 4). Specifically, P2 who had intact superior temporal lobe areas following ATL disconnection (Suppl. Fig. 2A), exhibited magnified functional connectivity that appears driven by these areas (Fig. 4). By contrast, P1 whose ATL disconnection procedure affected these anterior superior temporal lobe areas (Suppl. Fig. 2A) instead exhibited magnified interconnectivity that appears driven by the IFG (Fig. 4).

## DISCUSSION

This study presents unique intracranial data from the semantic neural network obtained with site-specific neurophysiological recordings before and within tens of minutes after a neurosurgical procedure for intractable epilepsy treatment that required ATL disconnection. In both participants, removal of the ATL resulted in robust alterations in neural signaling between IFG language sites and temporal lobe auditory speech processing sites distant to the surgical site (Fig. 1). Complete immediate compensation^17,18,20–22,24^ can be ruled out by the results, as can solely uncoordinated disruption to the neurophysiological system. Instead, the results, informed by the predictive coding framework, provide causal evidence in support of both the *diaschisis* and *incomplete functional compensation* hypotheses, providing direct causal evidence that the left hemisphere ATL has a vital role in speech processing and expectancy. The IFG, with its established role in semantic and syntactic language functions^8,9,11^, could coordinate possible compensation after the loss of the left ATL, together with the right ATL and speech-processing sites in the STG and HG. However, the intact areas, not directly physically affected by the ATL disconnection procedure, were unable to completely compensate for the impact on the network during this acute phase after the loss of the ATL.

Before ATL disconnection, the ATL showed hub-like structural and functional connectivity with the other nodes, consistent with its proposed centrality in the *hub-and-spokes* model^6,8^. Bottom-up evoked responses to physical acoustical features, namely voice onset time (VOT)^41,42^ were largely unaffected by ATL disconnection, while speech predictions and processes were widely disrupted in the recorded fronto-temporal network. The predictive coding framework^27,28^ anticipated both the form of alterations and possible compensation in the case of a loss of ATL prediction-related signals. The framework anticipated both the impact on semantic expectancy signals and the abnormally magnified gamma-band responses in sensory cortex (Fig. 2). Key evidence for attempted compensation^10,30^ was observed as increases in effective interconnectivity and a change in hub centrality in the fronto-temporal network post-disconnection. This included focal increases in functional connectivity between the fronto-temporal sites capable of compensating for a loss of speech-related and expectancy signaling in the neural network (Figs. 3-4). However, the intact regions and their functional interactions in this acute phase did not appear capable of recovering the depleted neurophysiological incongruency effects in HG after ATL disconnection (Fig. 1C). In addition, even two months post-surgery, we observed a lasting behavioral impact on the two patients’ abilities to integrate speech semantic and perceptual information (Fig. 1E). An intriguing observation, in terms of the predictive coding framework, was how consistently the disruption in interconnectivity between frontal and auditory sites affected the theta frequency band. Theta oscillations have a more ambiguous role in the predictive coding framework^43^, having been associated with both feedforward (entrainment to the speech syllabic rate^44,45^, cognitive sampling at a theta rhythm^46,47^) and feedback processes^48^. The neurophysiological results broadly support predictive coding and related frameworks, with testable extensions for the role of theta oscillations.

The acute alterations demonstrate the remarkable rapid capability of a neural network to respond to the loss of one of its hubs. The findings also provide important insights into the language connectome immediately before and after surgery affecting the left ATL, underscoring the limitations of cognitive models that either stipulate an irrecoverable functional role of the ATL or the possibility of complete immediate neurophysiological compensation by the rest of the network. The observations indicate that the left ATL is likely to act in two key ways: 1) as a higher-level site in a constructive hierarchy for semantic processing and predictions involving auditory cortex; and, 2) in interaction with language-critical areas in prefrontal cortex which have a proposed role in semantic control^49^.

Thereby, this study provides direct evidence for the vital role of the left ATL in the semantic network and validates predictive coding and related frameworks^50,51^, which can now be used to model acute neural network impact and compensation with greater confidence. These recordings —as some of the most immediate perioperative intracranial neurophysiological responses attainable from the human brain— also show remarkable neural network adaptability and rapid capability for compensation after the loss of a neural hub. Given the rarity of such ATL disconnection procedures, it would be important for future studies to establish direct correspondence source localized EEG^51–53^ or fMRI data^54,55^ in the same patients. Validated non-invasive imaging would then help to advance insights into neural network impact and compensation in larger patient cohorts throughout their extended recovery. We speculate that the form of rapid compensation seen here may engage early gene expression and adaptive brain plasticity mechanisms known to occur in tens of minutes^56,56^. The observed site-specific alterations could guide human patient or animal model research to further understand the cascade of compensatory and neurophysiological reorganization mechanisms on neural networks following neurological impact or treatment.

## METHODS

### Participants and ethical approvals

The research protocols were approved by the Institutional Review Board at the University of Iowa. Written informed consent was obtained from both participants. Participation in the research protocols was voluntary and did not interfere with the participants’ clinical treatment. The participants had the right to revoke consent at any time without affecting their clinical evaluation and treatment. Participant recruitment and research were conducted in accordance with current ethical principles and practices for conducting invasive intracranial research experiments in human patients^57^. This included maintaining the integrity of clinical care and space and ensuring the voluntariness of participation in research protocols at every stage. The research was carried out in one of the operating rooms at the University of Iowa Hospitals & Clinics during surgery for electrode removal and seizure focus resection.

### ATL neurosurgical disconnection procedure

We recorded data from the left hemispheres of 2 neurosurgical patients (P1: 63-year-old woman, P2: 32-year-old man). Both were native English speakers, right-handed and left hemisphere language dominant based on the Wada test. They had been diagnosed with medically refractory epilepsy and were undergoing surgery to remove seizure foci during acute electrocorticography (ECoG) monitoring in the operating room (OR). The surgery required the participants to be awake and able to interact with the clinical team for at least two periods of clinical motor speech and naming area mapping using electrical stimulation in order to minimize post-surgical morbidity^58^ (Suppl. Table 2).

An ATL disconnection procedure was performed for neurosurgical treatment, whereby to access the surgical treatment site a portion of ATL tissue is left intact within the middle cranial fossa and remains functionally disconnected from the remainder of the temporal cortex (transecting both gray and white matter)^59–61^. P1 participated in a two-step surgical procedure starting with the disconnection of the ATL. This made it possible for the clinical team to access the deeper epileptogenic foci structures for resection involving the hippocampus and parts of the amygdala, the second step in their procedure (Suppl. Fig. 2 shows the tissue removed during both steps). For P2, the procedure only required the first step, disconnection of the ATL to resect the epileptogenic cavernoma located in the temporal pole (Fig. 1B; Suppl. Fig. 2B). For both participants, we recorded data as they were awake before and after the ATL disconnection. The recording contact coverage is shown in Fig. 1C. This included temporal pole contacts, which only provided viable recordings before the ATL disconnection.

### Speech stimuli and behavioral task

The semantic prediction task, tested approximately a month before and 2 months after the surgical procedures, involved listening to sentences that primed the perception of a final target word (Fig. 1E), based on ^37^. It included naturally produced sentences to semantically bias the listener to expect a target word beginning with either a /b/ or /p/ phoneme at the final position in the sentence. The target word was manipulated along a VOT continuum with 6 steps (0 ms to 40 ms) to move evenly from the perception of /b/ as in ‘bark’ (short VOTs) to /p/ as in ‘park’ (long VOTs). Both endpoints corresponded to a real word, creating naturalistic conditions where the biasing sentence and the target word corresponded to the listener’s expectation or expectations were (congruent condition) or were not met (incongruent condition). Sentence primes were counterbalanced with target words from each step of the VOT continuum (Fig. 1E).

During the perioperative behavioral testing, the participants performed an extended version of the task. They were seated in an electrically shielded, sound attenuating room. During the semantic predictions task, 7 different word pairs were used (bill/pill, bad/pad, bath/path, back/pack, bark/park, bowl/pole, beach/peach), amounting to 504 experimental trials, not including 126 filler and 63 catch trials. Their task was to press a button that corresponded to the letter the target word started with (/b/ or /p/). The *filler trials* were sentences that fully predicted the target words, with no manipulation of the target word (e.g., “The walls need another coat of <u>paint</u>”, where *paint* is the target word which has no /b/ counterpart). The *catch trials* consisted of sentences where after hearing the sentence, the target word was presented visually on a screen along with another written word to choose from, in this case their task was to choose the words that best completed the sentences. These could either be fully predictive sentences as in the case of the filler trials, or fully congruent sentences taken from the main experiment. Altogether, the entire semantic predictions task and filler/catch trials consisted of 693 trials and took about 75 minutes to complete. Before the surgery, this task was also used to determine which 4 sentences had the biggest effect to focus on during the shorter data collection period possible in the operating room.

The phoneme identification responses on the semantic prediction task were statistically tested using a logistic mixed effects model with the three within-participant factors: VOT (6 levels: 1 to 6, centered), Semantic bias (2 levels: /b/- and /p/-bias, +/-0.5), and Disconnection (2 levels: 1 month before, and 2 months after the surgical procedure, +/-0.5). These were each entered as main effects along with the two-way interactions with disconnection. As these analyses were conducted individually for each subject, the only random factor was Word pair (7 pairs), and inferences allow us to generalize to the subject’s own behavior, not to the population. Following ^62^, we used the maximal random effects structure for both participants. Covariance terms among the random slopes were dropped to facilitate convergence. This led to the model given in (1), stated in R’s *lmer*() notation (see Supplementary Table 1 for complete results):

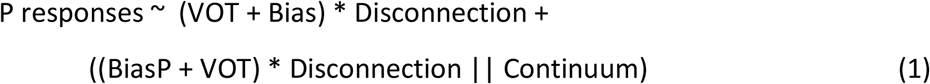

During the recordings in the operating room, the participants performed a passive version of the semantic predictions task to elicit neurophysiological activity corresponding to what they would have experienced during the pre- and post-operative testing. Stimulus delivery was controlled by the Psychophysics Toolbox^63,64^ running in MATLAB 2018b (The MathWorks, Inc., Natick, MA). Four different word pairs were used with three different preceding sentences (12 different sentences altogether) that biased listeners to expect one endpoint (/b/ or /p/) over the other, totaling 288 experimental trials. An additional 72 filler trials were also included, consisting of sentences that fully predicted their target word, to ensure that most trials fulfilled semantic expectations. Stimuli were presented at a comfortable level through a speaker placed in front of the participants. Each recording session was divided into 4 blocks so that the experimenter could ensure the participant was awake and engaged with the task. For P2, to additionally ensure the participant was awake and attending to the sounds presented, they participated in a simple tone-detection task during the speech priming paradigm. They were asked to press a button they were holding in their dominant right hand whenever a 440 Hz, 250 ms long sine-wave auditory tone was presented (24 randomly occurring tone detection trials in between the speech priming task). The first recording session was done after the clinical recording electrodes were placed, but before the ATL disconnection procedure, completing in 25-30 minutes. After the ATL disconnection procedure, a second similarly timed recording session with the same task ensued.

### Electrode coverage and neurophysiological data collection

Clinical platinum-iridium electrodes (AdTech, Racine, WI) were used to monitor interictal activity. The plans for placing ECoG electrodes for acute intraoperative clinical monitoring were formulated during a preoperative multidisciplinary epilepsy management conference. Four contact electrode strips were placed on the temporal pole, under the temporal lobe, and over the lateral surface of the superior temporal lobe. An 8-contact depth electrode was placed in HG using Stealth navigation (Medtronic). Two subgaleal electrodes were used as ground and reference. A 64- or 96-contact electrode grid (P2 and P1, respectively) was placed over the frontal lobe, containing reference and ground contacts. All contacts except the frontally placed grid remained in place for the entire recording time. The location of the frontal grid post-disconnection was checked and any displacement of recording contacts in relation to their pre-disconnection location was corrected. The positions of the frontal grids, and the STG electrode were confirmed after the surgery with the help of intraoperative photographs or in the case of HG using ultrasound^65^ to reconstruct the recording channels’ anatomical locations. The temporal pole and the subtemporal area locations could not be confirmed this way because they were occluded by the skull and are thus only approximated in location (Fig. 1C). The safety and clinical treatment utility of having the electrode coverage, including supratemporal HG electrodes has been previously demonstrated^66,67^. For instance, the pattern of ictal spread of seizure activity to the supratemporal plane is highly predictive of subsequent seizure-free outcome rates, and when high levels of interictal activity across the HG supratemporal plane electrode channels are observed, this is considered in the clinical resection plan.

### Structural and diffusion-weighted MRI

About a month before and approximately 2 months after the surgery, we obtained T1-weighted structural MRI scans (fast spoiled gradient-echo sequence, 1×1×0.8 mm resolution, TR=8.508, TE=3.288, flip angle=12^°^, 32 slices) and 64 direction diffusion-weighted MRI (1×1×2 mm resolution, TR=10332, TE=88.9, flip angle=90^°^) using a Siemens 3T Discovery MR750w scanner. Network construction methods broadly follow those described previously^51,68^. Preoperative T1-weighted MRI scans were processed using FreeSurfer version 6.0 using the *recon-all* tool to generate cortical regions-of-interest (ROI)^69^. The default parcellation scheme from FreeSurfer (the Desikan-Killiany atlas^69,70^) was used which contains 82 anatomical cortical and subcortical ROIs (e.g.,^71,72^). These regions were also combined to generate a gray matter ribbon using FSLmaths (later used as a seed region for surface-based tractography). Masks covering resected tissue were generated by first linearly registering (6 degrees of freedom) the post-operative T1-weighted MRI to the pre-operative T1-weighted MRI using FSL FLIRT. Then, using FSLview, we manually defined the resected tissue in pre-operative T1w space by overlaying the scans. Care was taken to avoid areas known not to be resected (see ^68^). Suppl. Fig. 2A shows the areas impacted by the disconnection of the left ATL in both participants.

Susceptibility distortions on preoperative diffusion-weighted MRI data were corrected using the SynB0 DisCo tool in conjunction with the FSL TOPUP tool^73,74^. The FSL tool ‘EDDY’ was used to correct for motion and eddy current distortions with the bvecs rotated appropriately^75^. The diffusion data were then reconstructed using generalized q-sampling imaging with a diffusion sampling length ratio of 1.25^76^. A deterministic fiber tracking algorithm was then used^77^. Tractography seeds were placed in the gray matter ribbon and deep brain structures by registering the preoperative *aparc+aseg* file generated by FreeSurfer to the preoperative diffusion scan (rigid body using *orig*.*mgz* DSI Studio). Tractography parameters were as follows: The anisotropy threshold was randomly selected. The angular threshold was 60 degrees. The step size was randomly selected from 0.5 voxel to 1.5 voxels. The fiber trajectories were smoothed by averaging the propagation direction with a percentage of the previous direction. The percentage was randomly selected from 0% to 95%. Tracks with length shorter than 15 mm or longer than 150 mm were discarded. A total of 5,000,000 tracts were calculated. Preoperative structural connectivity networks were then calculated as the number of streamlines ending between each region pair. To compute expected postoperative networks, we removed all tracts which pass into the mask voxels and recomputed the network. The difference between the two networks shows the percentage change in connectivity which we calculated as post- vs pre-disconnection. Networks were visualized using brainnetviewer^78^ (Fig. 2D).

### Intracranial ECoG acquisition and analysis

The ECoG data were sampled at 2000 Hz and filtered (0.7–800 Hz bandpass, 12 dB/octave roll off) using Tucker Davis Technologies system (TDT, Alachua, FL), then downsampled to 1000 Hz, denoised using demodulated band transform (DBT^79^; bandwidth = 0.25). Singular value decomposition was used to discard the first principal component based on the covariance matrix calculated from the highpass filtered (300 Hz) data. Event-related spectral perturbations were calculated with EEGlab^80^, epoched around the target word onset by log-transforming the power for each frequency and normalizing relative to the baseline (mean power in the pre-stimulus reference interval -150 to 0 ms before target word onset). The frequency band-related activity was averaged across all trials. The responses measured in each of the frequency bands (theta, alpha, beta, low and high gamma) were statistically compared using cluster-based non-parametric permutation testing, as described in ^81^, testing the contrast between the pre- and post-disconnection states (10,000 permutations, *p* < 0.05). We excluded any significant clusters shorter than 25 ms in duration.

Event-related potentials (ERPs) were calculated with respect to the semantic prediction context (i.e., whether the word was within a predicted congruent sentence context or an incongruent context). This allowed us to evaluate prediction mismatch-related neurophysiological responses (−150 to 800 ms from stimulus onset; baseline corrected and filtered off-line using a 0.5 to 50 Hz band-pass finite impulse response filter; Kaiser-windowed, Kaiser β=5.65, filter length=3,624 points). We calculated the ERPs to the congruent predicted condition using the 0 ms and 8 ms VOT target words for the /b/-bias sentences, and 32 ms and 40 ms VOT target words for the /p/-bias sentences. These were contrasted with the incongruent condition (e.g., prediction error) when the sentence created a bias towards expecting a /p/ word and the target word started with a /b/, using the 32 ms and 40 ms VOT target words for /b/-bias sentences, and the 0 ms and 8 ms VOT target words for /p/-bias sentences. The comparison included 96 trials each, and we subtracted the incongruent from the congruent ERPs to obtain a congruence difference wave. To account for the VOT differences expected on the neurophysiological response signals (Fig. 2A), we matched the congruent vs. incongruent conditions by VOT, extracting incongruent target words with the shortest VOT from congruent words with the shortest VOT, etc. We then compared the difference between the recordings before and after ATL disconnection. Statistical analysis used the non-parametric cluster-based permutation test as described for the spectral perturbation effects above.

All the available channels in each participant and ROI that had a significant speech response were evaluated, calculated by comparing the baseline pre-stimulus neurophysiological activity with the activity during the target word. If the latter exceeded 2 standard deviations from the baseline variability for over 125 ms of the 500 ms target word length, the channel was identified as having a significant speech response. The significant difference between the pre- versus post-disconnection speech response was evaluated using the cluster-based permutation test described earlier (*p* < 0.05). This process amounted to using 4 channels for each participant in HG, 2 in TP, 3 in STG. The IFG channels were included in the analyses only if they were localized to the IFG pars opercularis or triangularis after correction for the location of the IFG recording grid post-disconnection. IFG recording contacts in these regions were also only included for further analysis if they showed a significant speech response. For P1, this included 24 recording contacts for pars triangularis for both pre- and post-disconnection, and for pars opercularis, 11 and 10 channels respectively for pre- and post-disconnection. For P2, IFG recordings from pars triangularis included 11 channels pre-, and 9 post-disconnection, and for pars opercularis, 8 significant channels before and 5 after disconnection.

### State-space conditional Granger causality

Spectrally resolved state-space conditional Granger causality (CGC) analysis was used to investigate the directional influence between brain regions involved in the classification of the target speech word. The method is multivariate and conditional, in the sense that simultaneous time series from a collection of electrodes are included to account for direct and indirect influences between contacts. ECoG recordings were downsampled to 100 Hz and sectioned into trials, 0 to 2 seconds relative to the presentation of the target word. Intuitively, CGC tests whether activity in a source area can predict subsequent activity on a target area better than the target area can predict directional activity, as should be the case if the source area modulates activity in the recipient area. Prior to spectral CGC analysis, the mean at individual time points across trials was subtracted from the single trial ECoG data and then scaled by the standard deviation^82^. CGC considers the predictive effect of all other contacts, which allows us to distinguish between direct influences of interest, and artifactual indirect influences.

The state-space approach aims to address several significant problems in applying standard vector autoregressive (VAR) based CGC models to ECoG recordings, which are related to downsampling and nonstationarities^83^. The state-space model also addresses a number of theoretical and practical problems related to spectral CGC estimation^84–87^. Spectral CGC was directly computed using Geweke’s^88,89^ formulations based on the estimated state-space innovations covariance matrix, cross-spectral densities, and transfer functions^84,86^. State variables can be reconstructed from the measured ECoG recordings but are not themselves measured during an experiment. For modeling directional influence in the brain, it is possible to directly express the interactions between different regional signal time series as a state-space model. The state-space model in innovations form is defined by:

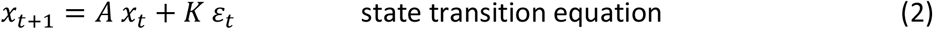

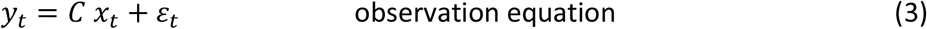

where *x*_*t*_ is an unobserved (latent) m-dimensional state vector, and *ε*_*t*_ is the vector of the innovations or prediction errors. The observed vector of time-series *Y*_*t*_ corresponds to the ECoG recordings from regions in the targeted network. The state transition matrix *A*, observation matrix *C* and the steady-state Kalman gain matrix *K* are estimated using a subspace method. Subspace methods are optimal for state-space model parameter estimation, especially for high-order multivariable systems^90^. The order of the state-space model was determined by the number of principal angles that differ from *π*/2^91^ when estimating the data-driven state-space model by the subspace method. Principal angles have been proven to be related to the singular values in singular value decomposition^91^. The order of the state-space model was 15, which corresponds to the vector size *x*_*t*_. Time series recorded from individual contacts were clustered together by averaging multiple simultaneous recordings to form ROIs. Post-stimulus time series from the two participants were concatenated as trials to combine the data.

To statistically evaluate the reliability of the connectivity results we used a phase-randomization surrogate data technique to construct an empirical null distribution^92–94^ representing chance influence between ROIs. This method consists of randomly shuffling the Fourier phases of each of the ECoG recordings, which generates uncorrelated data with preserved autocorrelation properties. The matrix of spectral CGC between four ROIs in HG, STG and IFG opercularis and triangularis was statistically evaluated as follows: 2,000 surrogates were generated and thresholded at *α* = 0.05 of the null distribution, values below which were subtracted from the observed spectral CGC and trimmed at zero. Finally, statistically significant pre- and post-disconnection spectra were plotted together for comparison. For both cases significant influence between ROIs was observed across the range of frequencies studied up to 50 Hz. To examine specific frequency band influences, the spectrum was segmented into four bands: theta (4-8 Hz), alpha (9-12 Hz), beta (13-30 Hz) and gamma (31-50 Hz). Average CGC magnitudes were computed for both directions between ROI pairs. Permutation tests were performed for each frequency band by constructing a null distribution of signed pre- and post-disconnection spectrally segmented differences following recalculation of spectral CGC after random assignment of time series to the two categories. To compare whether directional influences between ROIs change between pre- and post-disconnection, random permutation tests were performed on the absolute difference between spectral CGC across frequencies. Each post-stimulus trial time series was randomly assigned to either the pre- or post-disconnection category and spectral CGC recalculated 2,000 times. An empirical null distribution was calculated by summing the absolute CGC differences between pre- and post-disconnection across frequencies. The number of permuted surrogates that exceeded the measured difference was used to calculate *p*-values. False discovery rate (FDR) was controlled at the 0.05 level^95^ across ROI pairs.

### Neural speech representation patterns and phase amplitude coupling

Neural pattern similarity was calculated across recording contacts in response to the speech sounds. First, high gamma band (70-150 Hz) responses were averaged over the target word epochs (between 0 ms to 800 ms during the target word presentation period) for the shortest and the longest VOT. This amounted to 48 trials for the 0 ms /b/ and the 40 ms /p/ target words each. For this analysis, all temporal lobe recording channels were used, excluding the TP electrodes because they were not viable after disconnection and one STG electrode in the case of P1 which was also deemed unusable prior to analysis. Then correlation matrices using Spearman rank correlation were calculated between electrodes with respect to the target words split across two semantic sets determined by the shortest and longest VOTs. The neural pattern dissimilarity matrices were calculated (1-correlation coefficient) for the datasets recorded before and after the disconnection. Then, all four correlation matrices, excluding the diagonal and redundant matrix items, contributed to an overall dissimilarity matrix to assess the impact on the multi-contact pattern of responses to the target words before and after ATL disconnection. Permutation testing with 5,000 replicates was used. Here, surrogate Spearman correlation coefficients were computed with the neural pattern shuffled at random. The *p* value was taken to be the proportion of replicates where the surrogate values exceeded the observed value. Significance was determined by thresholding at α = 0.05 with Bonferroni correction.

We also assessed the impact of ATL disconnection on phase-amplitude-coupling (PAC) in response to the speech sounds, given that PAC is a feature of auditory cortical responses to speech^43,96^. We calculated PAC in response to the target words per recording site using a PAC modulation index (MI)^43^. The MI evaluated the strength of coupling between low-frequency phase (4–8 Hz) and high-frequency amplitude (40–100 Hz). The matrix of MI values was *z*-score thresholded (*z* > 3) and averaged across HG channels by participant (*N* = 4 for each participant). The MI matrix was computed during the entire stimulus duration (800 ms) for all pairs of amplitude (25–100 Hz in 5 Hz steps) and phase (4–8 Hz in 1 Hz steps). This was done by concatenating across trials to increase the signal-to-noise ratio needed to estimate cross-frequency coupling and detect coupling effects. The statistical significance threshold for MI values was determined using Bonferroni correction for multiple comparisons (*α* = 0.05, 90 comparisons of pixels in the MI matrix, corresponding to a corrected *p* < 0.05 and *z* score of 3.3 or higher). The comparison between the pre- and post-disconnection data was done by a Wilcoxon signed-rank test.

### Minimizing anesthetic and direct neurosurgical confounds

To optimize the quality of clinical ECoG recordings and minimize the effects of anesthesia, surgical anesthesia had been discontinued in both participants at least 20 minutes before the pre- and post-disconnection recording periods. Vital signs were similar and stable during both pre- and post-disconnection periods (e.g., heart rate and respiration: Suppl. Table 2). The participants were awake during the recordings, and we were able to obtain data in P2 on a tone-detection paradigm during the speech priming task, indicating that they were alert and similarly engaged during the pre- and post-disconnection recording periods. Immediate post-anesthesia drowsiness is expected, but the experimenter monitored both participants throughout the task and recording periods to ensure that they remained alert and similarly engaged throughout both pre- and post-disconnection. Both before and after ATL disconnection recordings were conducted after the craniotomy procedure and would be similarly affected by it. The ATL disconnection step in the surgical procedure can be expected to alter brain functions either through direct impact on neural elements, or as a result of edema formation within tissue adjacent to the surgical site^97–100^. Edema is an implausible cause of the observed effects because the intact auditory-frontal areas recorded from here post-disconnection were located several centimeters away from the surgical disconnection site. Also, the responses to the speech sounds are consistent with prior reports conducted in the awake state, in both form and amplitude^40,101,102^, and we observed little impact on auditory cortex VOT representations post-disconnection (Fig. 2A). Furthermore, whereas non-specific neurosurgical effects are expected to disrupt and minimize effective connectivity post-disconnection, specifically magnified speech responses or effective connectivity are difficult to explain by general neurosurgical effects alone. These effects are better explained by cognitive and predictive coding framework models, see text. These considerations suggest minimal impact of non-specific anesthesia or disruptive neurosurgical effects on the reported neurophysiological processes before and after ATL disconnection.

## ACKNOWLEDGEMENTS

Supported by Wellcome Trust (CIP: WT092606AIA; TDG: WT091681MA), National Institutes of Health USA (MAH: R01–DC04290; U01-NP103780) and European Research Council (CIP: ERC CoG, MECHIDENT). P.N.T. is supported by a UKRI Future Leaders Fellowship [MR/T04294X/1]. We thank Dr. R. Mueller and H. Chen for assistance with data collection, F. Balezeau for help with the dMRI processing, and T. Abel, M. Banks, M. Long, K.V. Nourski, H. Oya, B. Park and M. Steinschneider for discussion.

## DATA SHARING STATEMENT

The data to replicate the results of this study would be openly shared with its publication on the Open Science Framework: https://osf.io/arqp8/.

## AUTHOR CONTRIBUTIONS

Zsuzsanna Kocsis: Data collection, Conceptualization, Methodology, Software, Formal analysis, Visualization, Writing – original draft, review & editing. Rick L Jenison: Conceptualization, Methodology, Software, Formal analysis, Investigation, Visualization, Writing – original draft, review & editing. Thomas E Cope: Investigation, Writing – original draft, review & editing. Peter N Taylor: Software, Formal analysis, Investigation, Writing – original draft, review & editing. Ryan M Calmus: Visualization, Investigation, Writing – review & editing. Bob McMurray: Conceptualization, Methodology, Writing – review & editing. Ariane E Rhone: Conceptualization, Methodology, Writing – review & editing. McCall E Sarrett: Data collection, Software, Conceptualization, Methodology, Writing – review & editing. Yukiko Kikuchi: Software, Formal analysis, Investigation, Writing – review & editing. Phillip E Gander: Investigation, Writing – review & editing. Joel I Berger: Data collection, Formal analysis, Writing – review & editing. Christopher K Kovach: Investigation, Writing – review & editing. Inyong Choi: Investigation, Writing – review & editing. Jeremy D Greenlee: Investigation, Writing – review & editing. Hiroto Kawasaki: Investigation, Writing – review & editing. Tim D Griffiths: Conceptualization, Writing – review & editing, Supervision, Funding acquisition. Matthew A Howard III: Conceptualization, Writing – review & editing, Supervision, Project administration, Funding acquisition. Christopher I Petkov: Conceptualization, Methodology, Visualization, Investigation, Funding acquisition, Supervision, Writing – original draft, review & editing.

## SUPPLEMENTARY MATERIALS

**Suppl. Fig. 1:** Behavioral /b/ bias condition results

**Suppl. Fig. 2:** Resection impact on cortical tissue, connectivity matrix similarity across participants and P1 simulated changes to connectome only including TP impact

**Suppl. Fig. 3:** Temporal Pole (TP) pre-disconnection Conditional Granger Causality (CGC) results

**Suppl. Fig. 4:** Disconnection speech response impact on HG, IFG and STG across frequency bands

**Suppl. Fig. 5:** Speech predictability mismatch effects in HG, STG and IFG

**Suppl. Fig. 6:** Neural representational pattern and speech phase-amplitude (theta-gamma) coupling.

**Suppl. Fig. 7:** CGC results by congruent and incongruent conditions

**Suppl. Table 1:** Results of logistic mixed effects model for the behavioral task for each participant.

**Suppl. Table 2:** Physiological parameters during the pre- and post-disconnection recording periods

**Supplementary Figure 1.**
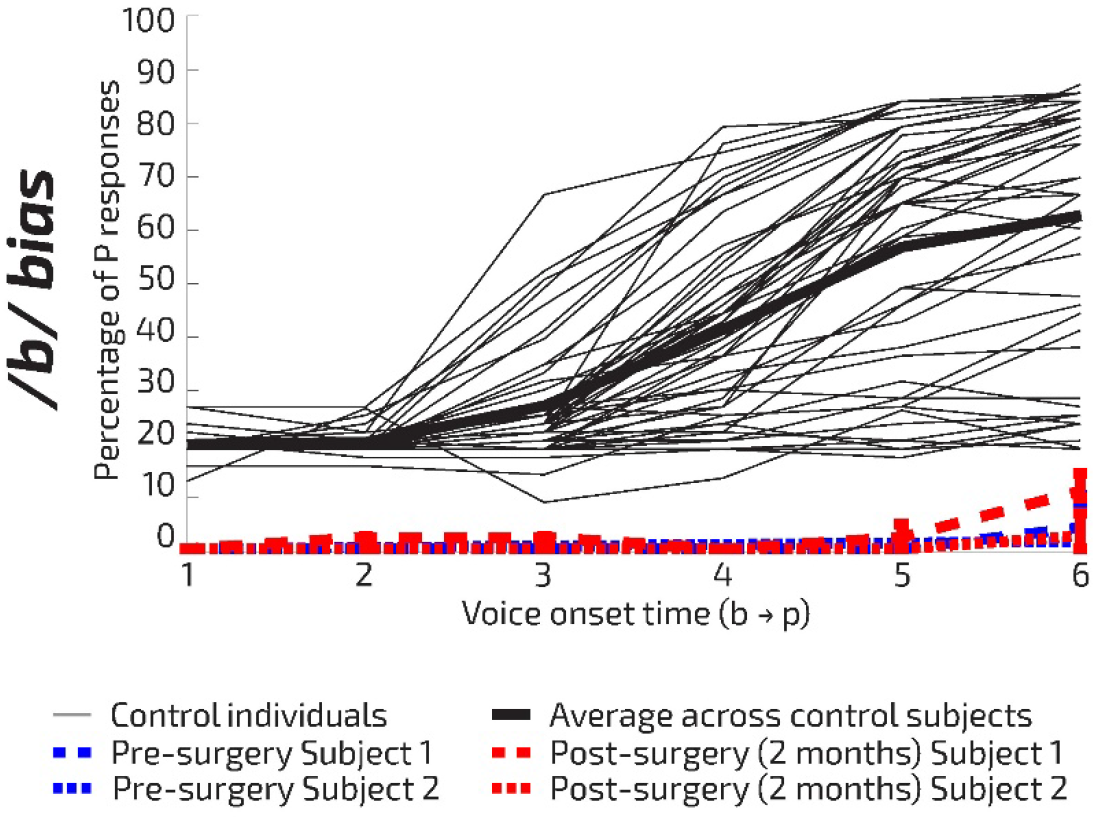
Behavioral /b/ bias condition results. Percentage of P responses to the /b/ bias words in relation to the VOTs (from /b/ to /p/ sounds) from the control participants and P1 and P2. The P1 and P2 data were obtained 2-6 weeks before and two months after their surgery. Format as in manuscript Fig. 1F. The /p/ bias responses (manuscript Fig. 1F) are well within the range of responses of the control participants’ data pre-disconnection and are substantially disrupted post-disconnection in both P1 and P2. The /b/ bias responses of both participants shown here are outliers both before and after the surgery. The combined /b/ and /p/ behavioral results for P1 post disconnection suggest that after the loss of the ATL, P1 adopts a strategy based primarily on the context of the preceding sentence. For P2, the combined /b/ and /p/ behavior after the ATL disconnection procedure suggests a different strategy, less based on the sentential context, but one unlikely to be solely a /b/ button response bias because both participants performed very well on the control trials that are interleaved in the experimental task that require both /p/ and /b/ button responses.

**Supplementary Figure 2.**
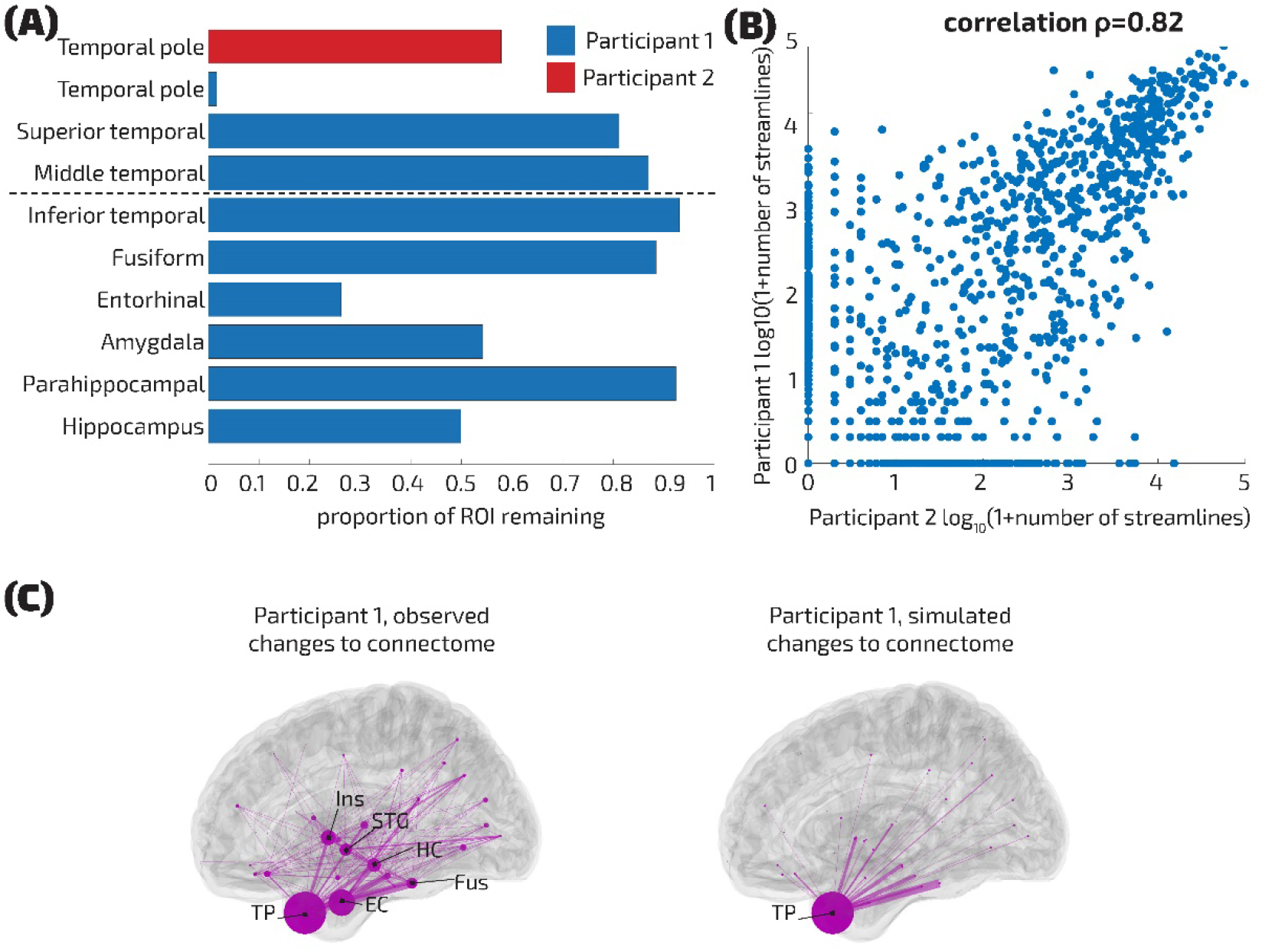
Resection impact on cortical tissue, connectivity matrix similarity across participants and P1 simulated changes to connectome only including TP impact. (A) The amount of the given region left after the entire surgical procedure based on the T1-weighted scans recorded before and two months after the disconnection of the left ATL for P1 and P2. P1’s entire resection procedure involved both the ATL disconnection and para/hippocampal resection (blue bars). The dashed line denotes the separation of the two steps of the surgical procedure: in step 1, the ATL is disconnected, and the superior and middle temporal gyri are also affected (after this step, our post-disconnection dataset was collected); in step 2, deeper structures are resected, the following brain regions are affected: inferior temporal and entorhinal cortex, fusiform and parahippocampal gyrus, amygdala and hippocampus. P2’s procedure only involved the ATL disconnection affecting the TP (red bar). (B) Similarity of connectivity matrices before disconnection. Shown is the comparison of P1 and P2’s log transformed number of streamlines between the nodes in their connectome, based on the dMRI images before disconnection. (C) Observed reduction (left) in region-region streamlines (proportional to line width) and hubness (degree centrality; proportional to node diameter) in P1 after ATL and para/hippocampal resection (TP: temporal pole; Ins: insula; STG: superior temporal gyrus; HC: hippocampus; EC: Entorhinal cortex; Fus: Fusiform gyrus). Also shown is the simulated structural impact of disconnecting only the TP node (right). The empirically observed impact includes the expected reduction in TP centrality seen in the simulation, and all edges disconnected within the simulation were also disconnected, featuring prominently in the observed impact. The simulated partial surgical impact is therefore consistent with the observed post-surgical impact, and plausibly representative of the status of the disconnection at the time of the functional recordings.

**Supplementary Figure 3.**
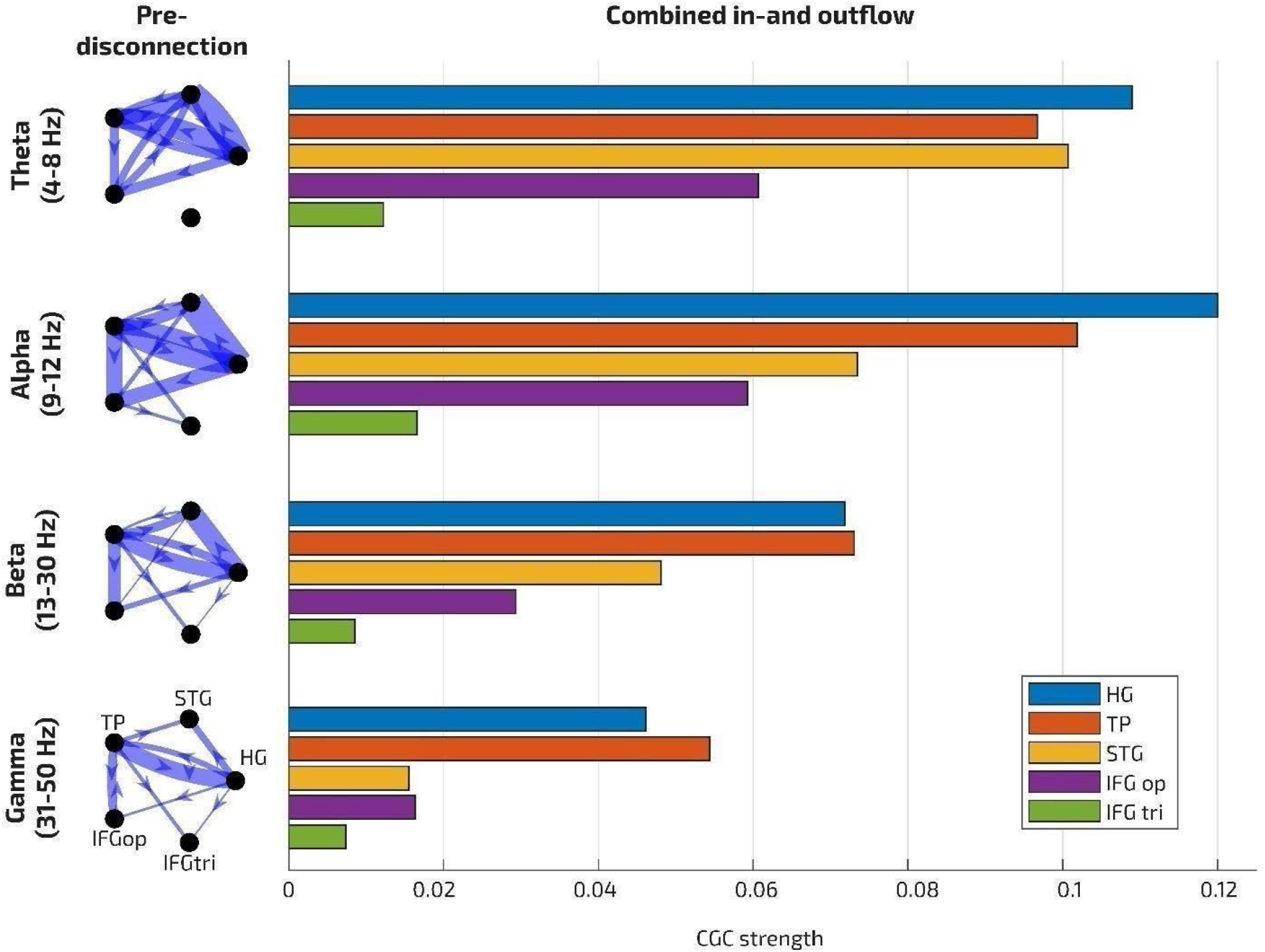
Temporal Pole pre-disconnection Conditional Granger Causality (CGC) results. Right: Directions of influence are shown from the regions of interest to recipient regions shown to be active during the target speech sound: HG, STG, TP and IFG pars triangularis and opercularis. Subthreshold (non-significant) regions of time-frequency CGC are masked (set to 0). Left: Combined in- and outward edge weights from and to each region in each frequency band, showing ‘hub-like’ activity in TP in higher (beta and gamma) and in HG in lower (theta and alpha) frequency bands. IFG p. triangularis of all the regions is least hub-like pre-disconnection.

**Supplementary Figure 4.**
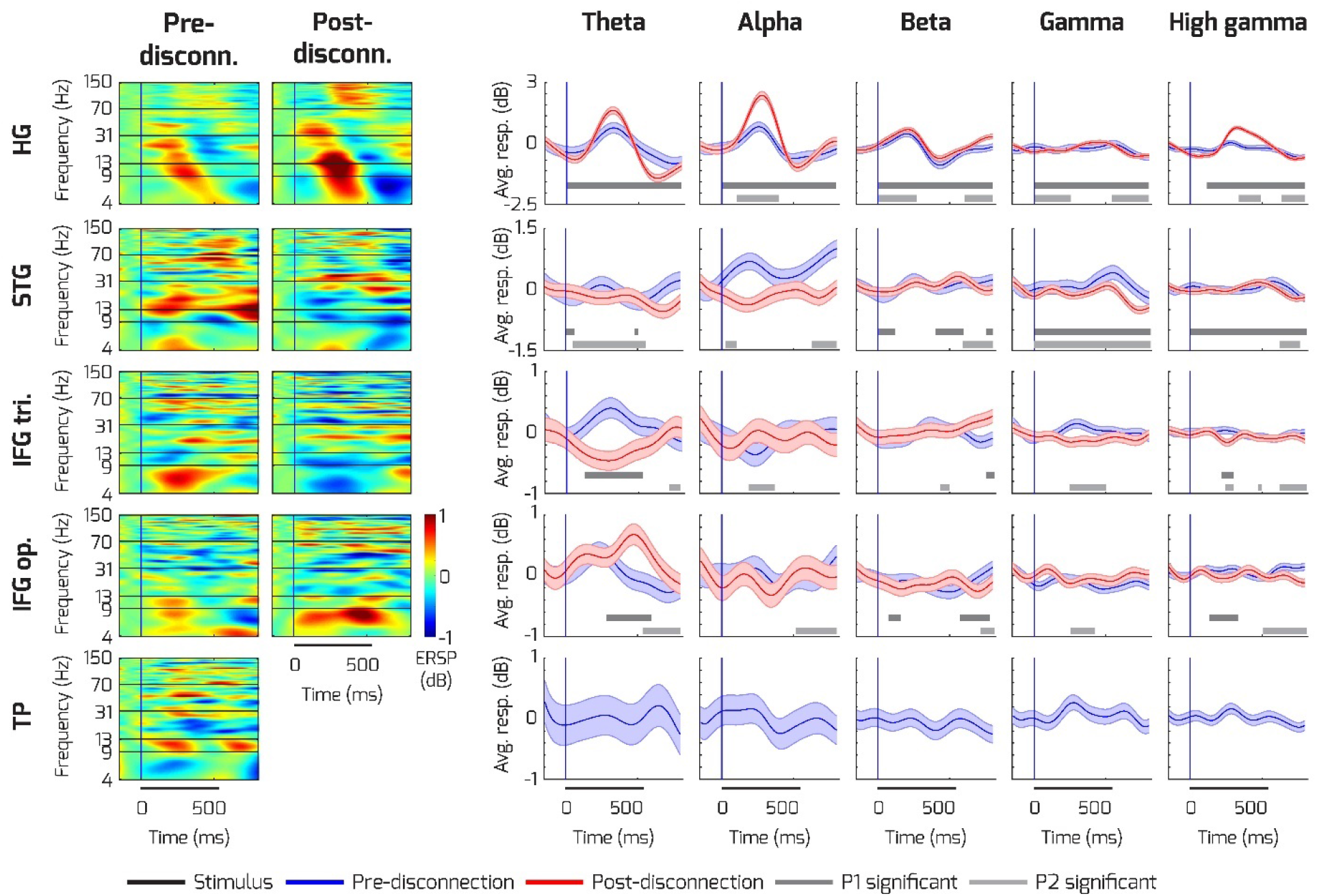
Disconnection speech response impact on HG, IFG, STG and TP across frequency bands. Format as in manuscript Fig. 2B. Left two columns: Event-related spectral perturbation responses to the target speech sound (stimulus onset indicated by black bar below plots) in HG, STG, IFG pars triangularis and opercularis and TP electrodes for both participants. TP contacts were only neurophysiologically viable pre-disconnection. Right: Averaged theta (4-8 Hz), alpha (9-12 Hz), beta (13-30 Hz), gamma (31-69 Hz) and high gamma (70-150 Hz) frequency band responses plotted together with the standard error of mean (SEM). Blue lines show the responses before, red lines show the responses after disconnection of the ATL. Gray bars indicate permutation tested significant differences post- vs pre-disconnection for P1 and P2, respectively (cluster-based permutation test p < 0.05 for at least 25 ms time windows). Unlike HG which shows substantial magnification of speech responses (also see manuscript Fig. 2B), STG shows largely disruption of speech representations post-disconnection as does IFG pars triangularis. IFG pars opercularis shows theta speech response enhancement after disconnection.

**Supplementary Figure 5.**
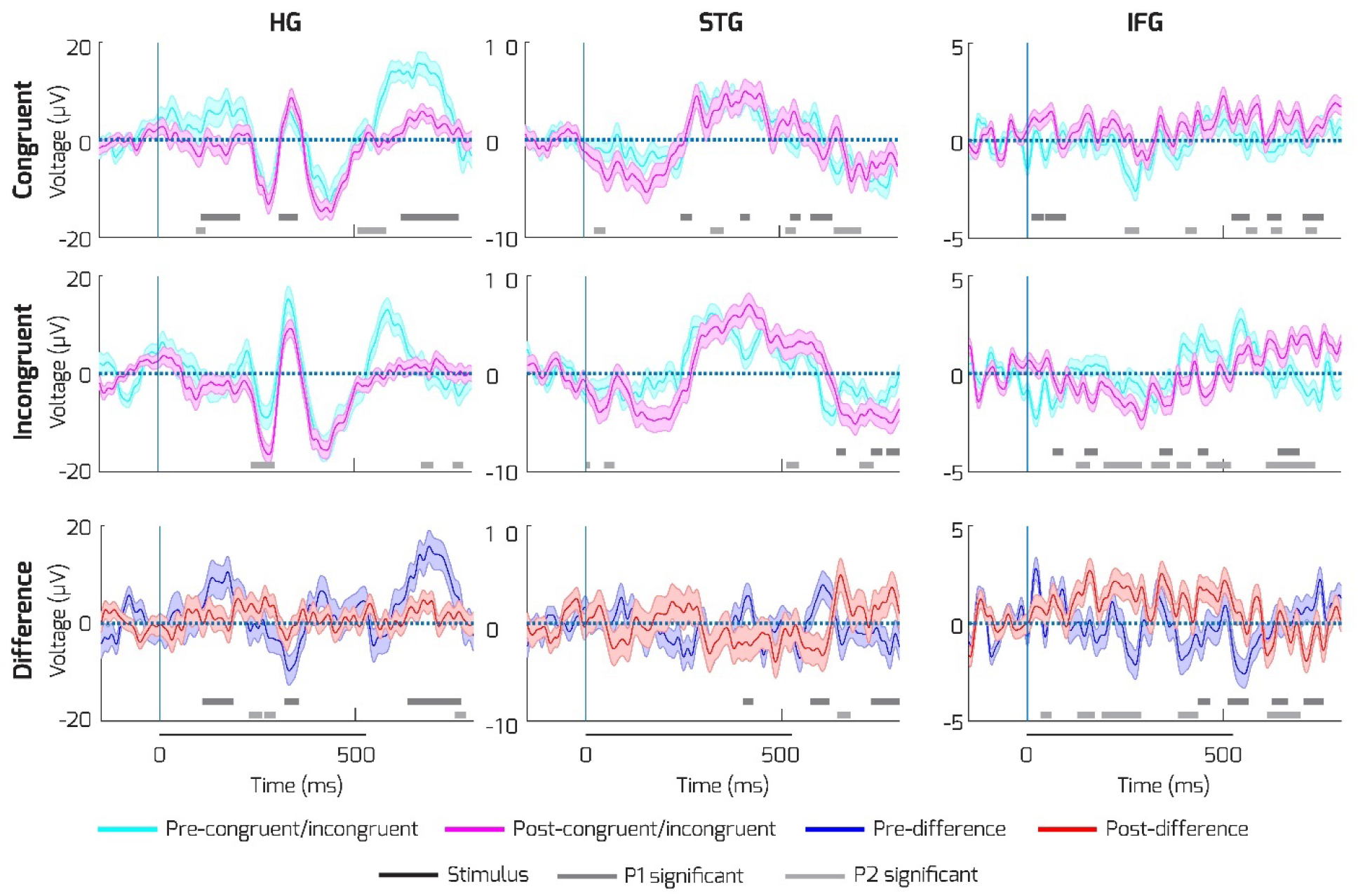
Speech predictability mismatch effects in HG, STG and IFG. Top row shows the responses to the congruent condition before (cyan) and after (magenta) the disconnection. The second row shows the same but for the incongruent condition. The third row shows the contrast between congruent and incongruent conditions before (blue) and after (red) the disconnection. The solid black line under the time axis depicts the average length of the target speech stimuli. The gray lines under the plots depict significantly different clusters for P1 and P2, respectively (Methods). Gray bars indicate permutation tested significant differences post- vs pre-disconnection for P1 and P2, respectively (cluster-based permutation test p < 0.05 for at least 25 ms time windows). Format as in manuscript Fig. 2C. HG congruency effects are strong in HG and disrupted post-disconnection (bottom left). Such effects are weaker in STG and IFG but also show signs of disruption post-disconnection.

**Supplementary Figure 6.**
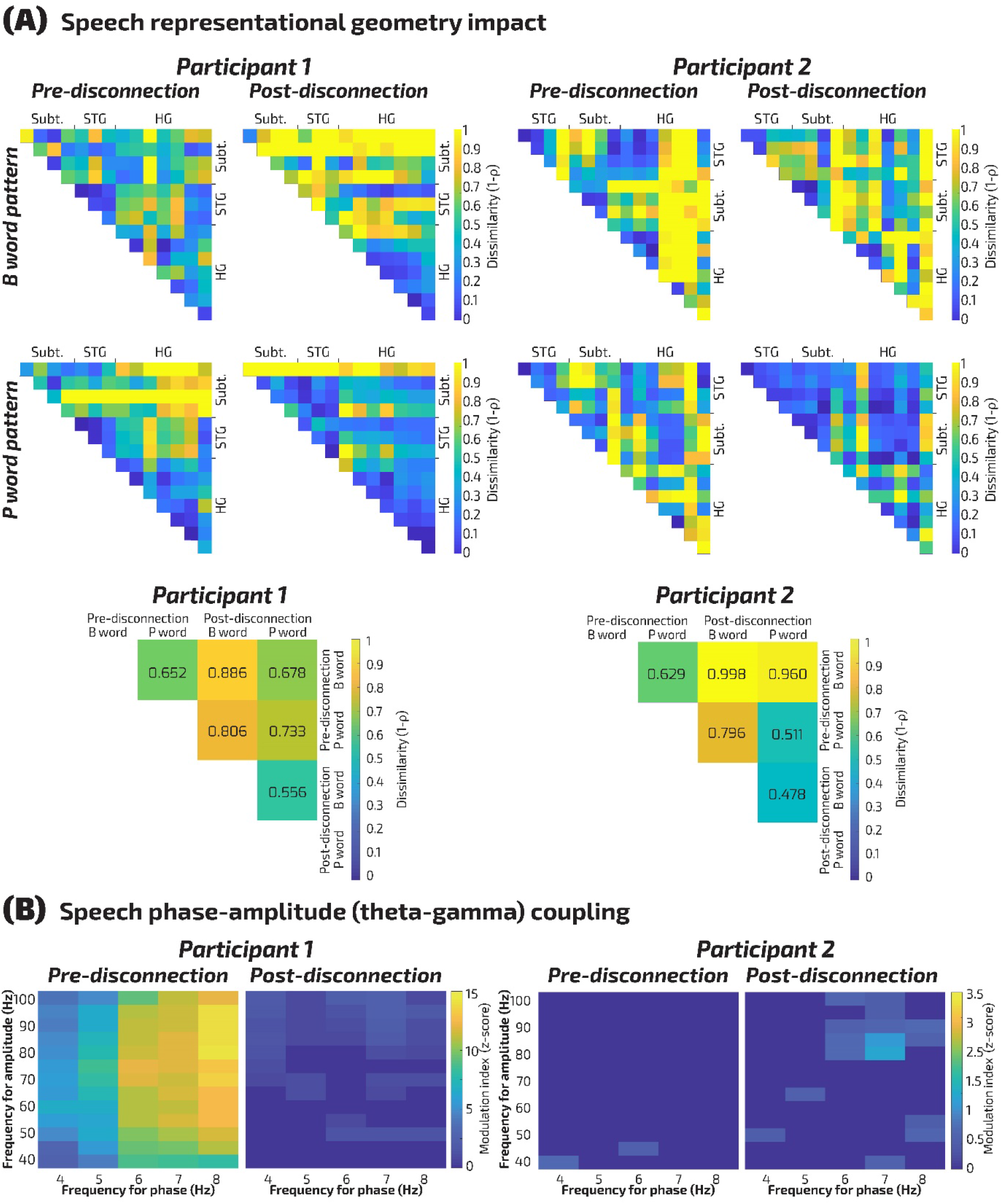
Neural representational pattern and speech phase-amplitude (theta-gamma) coupling. (A) Upper panel: 0 ms /b/ and 40 ms /p/ word neural representation patterns across all temporal lobe recordings contacts (HG, STG, Subt: subtemporal) shown as a dissimilarity matrix (1-Spearman *ρ*) for P1 and P2. Lower panel: Dissimilarity matrix summarizing the pattern change between pre- and post-disconnection in both participants. The neural response pattern to the /p/ words remained significantly similar post-disconnection, but the /b/ lost its similarity. We evaluated the ATL disconnection impact on neural response patterns to the speech sounds, cast as a multivariate pattern of responses across all of the temporal lobe contacts to the sets of /b/ and /p/ words. We tested the null hypothesis that the loss of the ATL would maintain the similarity of the representations pre- and post-disconnection. We found that the neural pattern results were consistent across both participants in the /b/ word pattern not being significantly similar after ATL disconnection (P1: *p* = 0.128; P2: p = 0.478). (B) Impact of ATL disconnection on Phase Amplitude Coupling (PAC) in response to the speech sounds. Left: Phase-amplitude (theta-gamma) coupling to the speech sounds was significant in P1 before disconnection and significantly disrupted post-disconnection (*p* < 0.001; Wilcoxon signed-rank test). Right: P2 did not show significant theta-gamma coupling to speech pre-disconnection thus the post-disconnection impact in P2 could not be determined. In both participants, the /p/ pattern remained similar pre-relative to post-disconnection (P1: *p* = 0.002; P2: p = 0.0002). Theta-gamma coupling was only evident pre-disconnection and significantly disrupted post-disconnection in P1.

**Supplementary Figure 7.**
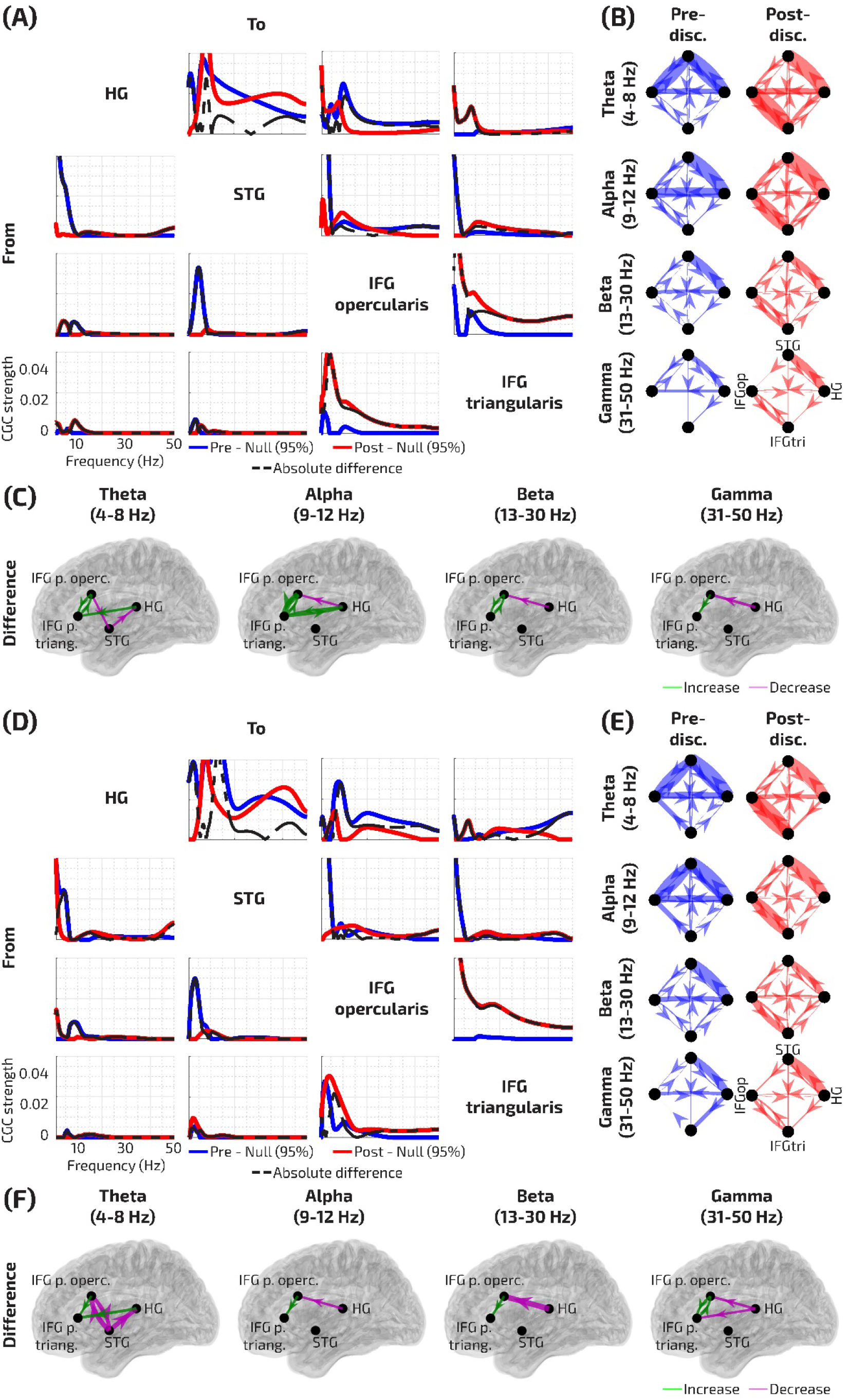
CGC results by congruent and incongruent conditions. Format as in manuscript Fig. 4. (A) Directions of influence are shown from regions of interest (rows) to recipient regions (columns) shown to be active during speech categorization: HG, STG, and IFG pars triangularis and opercularis. Only statistically significant effects are shown for all spectra, with the phase-randomized null distribution 95th% subtracted for each of the pre- (blue) and post-disconnection (red) conditions (B) Strong dynamic directional influences are observed between the state-space modeled time-series and IFG in the congruent word condition. (C) The difference between pre- and post-disconnection directional influences across the brain regions in the congruent word condition. (D) Same as (A) but for the incongruent condition. Only statistically significant effects are shown for all spectra. (E) Strong dynamic directional influences are observed between the state-space modeled time-series and IFG in the incongruent word condition. (F) The difference between pre- and post-disconnection directional influences across the brain regions in the incongruent word condition. The broad band magnification of effective connectivity between the two IFG subregions post-disconnection is evident for both the congruent and incongruent conditions. We observe that congruency-related effects were enhanced in interaction between the IFG and auditory cortex post-disconnection particularly in the alpha band and enhanced IFG and HG interactions in theta oscillations.

**Supplementary Table 1.** Results of logistic mixed effects model for the behavioral task for each participant.

|  | **P1** |  |  |  | **P2** |  |  |  |
| --- | --- | --- | --- | --- | --- | --- | --- | --- |
| **Factor** | **B** | **SE** | **Z** | **p** | **B** | **SE** | **Z** | **p** |
| *VOT* | 3.77 | 0.56 | 6.71 | <0.0001 | 3.32 | 0.75 | 4.45 | <0.0001 |
| *Bias* | 5.95 | 0.87 | 6.83 | <0.0001 | 4.60 | 0.66 | 6.93 | <0.0001 |
| *Disconnection* | 2.06 | 0.60 | 3.44 | 0.0006 | -1.37 | 0.70 | -1.95 | 0.0514 |
| *Disconnection* × *VOT* | -3.40 | 1.23 | -2.77 | 0.0056 | 0.93 | 1.01 | 0.92 | 0.36 |
| *Disconnection* × *Bias* | -0.71 | 1.19 | -0.59 | 0.55 | -1.30 | 1.29 | -1.01 | 0.31 |

**Supplementary Table 2.** Vital signs for the two participants during the intracranial recording periods and anesthesia drug stoppage before the awake recordings.

|  | **Participant 1** |  | **Participant 2** |  |
| --- | --- | --- | --- | --- |
|  | **Pre-disconnection** | **Post-disconnection** | **Pre-disconnection** | **Post-disconnection** |
| *Time of recording* | 11:34 – 12:03 | 14:48 – 15:15 | 12:38 – 13:07 | 15:03 – 15:34 |
| *Dexmedetomidine stopped* | 10:59 – 13:24<br>(35 minutes before testing period) | 14:25 – 15:24<br>(23 minutes before testing period) | 11:09 – 13:27<br>(37 minutes before testing period) | 14:40 – 15:34<br>(23 minutes before testing period) |
| *Heart rate (avg; +/- SD) during recording period* | 59.19 (1.28) | 66.93 (3.25) | 58 (3.91) | 55.94 (1.77) |
| *Respiration rate (average/minute; +/- SD) during the recording period* | 16.25 (0.51) | 18.59 (1.02) | 15.71 (2.6) | 13.45 (4.12) |

